# Single-Shot Optical Projection tomography for high-speed volumetric imaging

**DOI:** 10.1101/2021.10.15.464407

**Authors:** Connor Darling, Samuel P.X. Davis, Sunil Kumar, Paul M.W. French, James McGinty

## Abstract

We present a single-shot adaptation of Optical Projection Tomography (OPT) for high-speed volumetric snapshot imaging of dynamic mesoscopic samples. Conventional OPT has been applied to *in vivo* imaging of animal models such as *D. rerio* but the sequential acquisition of projection images required for volumetric reconstruction typically requires samples to be immobilised during the acquisition of an OPT data set. We present a proof-of-principle system capable of single-shot imaging of a 1 mm diameter volume, demonstrating camera-limited rates of up to 62.5 volumes/second, which we have applied to 3D imaging of a freely-swimming zebrafish embryo. This is achieved by recording 8 projection views simultaneously on 4 low-cost CMOS cameras. With no stage required to rotate the sample, this single-shot OPT system can be implemented with a component cost of under £5,000. The system design can be adapted to different sized fields of view and may be applied to a broad range of dynamic samples, including fluid dynamics.

## Introduction

Optical 3D imaging of intact samples at mesoscopic (mm-cm) scales has proved a valuable capability in biomedicine, enabling 3D structures to be observed in their native context. Furthermore, the use of non-ionising radiation and the wealth of fluorescent markers in the visible region enables specific molecular imaging, including *in vivo* studies of small transparent organisms. Successful volumetric fluorescence imaging techniques include optical projection tomography (OPT)^1^ and selective plane illumination microscopy (SPIM)^2^ also described as light sheet fluorescence microscopy (LSFM)^3,4^ and these have been applied to a range of *in vivo* studies^5–9^. However, it is usually necessary to immobilise live samples since the 3D image data set is constructed from sequential image acquisitions over timescales during which the sample should remain stationary. Organisms such as zebrafish embryos are therefore typically imaged under anaesthetic and often suspended in a gel such as agarose^5,6,10,11^. The required imaging time (and therefore the physiological stress on the organism^12^) can be minimised by increasing the rate of the 3D image data acquisition^13^ and/or by reducing the total number of images required for 3D reconstruction, e.g. using compressive sensing techniques^5^. Nevertheless, all 3D imaging techniques that rely on sequential image acquisition can be prone to motion artefacts and, for highly dynamic samples, such as fluids mixing, 3D imaging flow cytometry assays and small organisms in motion, it would be desirable to acquire the whole 3D image data set in single shots at the maximum frame rate of the camera. To date, single-shot 3D volumetric imaging has only been demonstrated with holographic techniques^14^ (so not applicable to fluorescence imaging) or parallax-based methods, such as stereo microscopy or plenoptic^15^ imaging, where the 3D image information is limited in angular coverage.

Optical projection tomography (OPT)^1^ is the optical analogue of X-ray computed tomography for which a series of widefield projection views through a sample are recorded as it is rotated through 360°. The volumetric image is then reconstructed from the set of projection measurements, typically using filtered back projection (FBP)^1,16^. Since the key components of an OPT system are a motorised rotation stage, a camera, a telecentric imaging system and a light source (which can be an LED), OPT is low-cost compared to most other volumetric imaging modalities. As wide-field imaging can be easily scaled to large (cm) fields of view using an optical imaging system with appropriate magnification, OPT has been applied to cm scale chemically cleared^17^ tissue samples for preclinical studies, e.g. whole mouse brains^18^, mouse pancreas^19^, adult zebrafish^20^, and clinical pathology^21^. For such *ex vivo* applications, total image acquisition time is not a critical parameter and highly sampled fluorescence OPT projection data sets can typically be acquired on a timescale of minutes – noting that a fully Nyquist sampled OPT data set requires projection images acquired at π*N* angles, where *N* is the number of resolution elements across the projection image.

OPT is also applicable to the imaging of transparent (weakly scattering) *in vivo* models (e.g. *C. elegans*^22,23^ and *D. rerio*^5,8,10^ and can be minimally toxic through the use of non-ionising radiation. However, with OPT data acquisitions typically requiring several minutes per colour channel, live samples may be compromised by phototoxicity and the stress of being maintain under anaesthesia, limiting the applicability for longitudinal studies. While the total OPT data acquisition time can be reduced by acquiring sparse (angularly under-sampled) OPT data sets on shorter timescales (~ 10 seconds) and using compressed sensing techniques^5,24^ or machine learning^25^ to minimise “streak artefacts” in the reconstructed volume, any OPT approach that still requires the sequential acquisition of multiple projection images may be compromised by samples moving on the timescale of sequential OPT image acquisitions.

We note that full volumetric single-shot imaging could be realised using OPT implemented with multiple cameras each acquiring a separate projection image in parallel. This would be prohibitively expensive to implement in a fully sampled OPT system (which typically requires several hundred projections) and the size of the resulting data sets would be prohibitively large for high speed volumetric imaging. By utilising compressive sensing, however, it is possible to reduce the number of projections (and therefore cameras) required and thereby reduce the size of the acquired data sets. Here we present the first demonstration of single shot volumetric imaging using compressing sensing OPT that is capable of imaging ~ 1 mm diameter volumes at the camera frame rate, achieved by recording 8 projection views simultaneously using 4 cameras. We demonstrate the 3D volumetric imaging of a free swimming zebrafish embryo at 62.5 volumes/second (limited by the frame rate of the cameras currently available to us). As well as enabling imaging of dynamic samples, single-shot OPT (ss-OPT) eliminates the need for sample rotation and so could also be applied to *in situ* long timescale imaging e.g. for application in developmental biology. Using recently available low-cost CMOS cameras, which feature a global shutter and the ability to receive a hardware trigger, the total component cost of the single-shot OPT system presented here is below £ 5,000.

## Methods

### Angular multiplexing OPT

In order to record two projection views on each camera, an angular multiplexing OPT^10,26^ optical setup was implemented in a novel compact configuration. This approach takes advantage of “excess” numerical aperture (NA) of the imaging objective lens, noting that the requirement for parallel ray projections for OPT, which corresponds to the imaging depth of field being greater than half the sample diameter, places an upper limit on the NA of the imaging system. This limited NA is typically imposed using an aperture in the Fourier plane of the imaging objective lens. In our OPT system implemented with 4x magnification microscope objectives (PF4X-INF, AmScope), this maximum NA required by the OPT system is much less than the full NA (0.13) available from these objective lenses. It is therefore possible to make more use of the angular collection of the lenses by placing more than one limiting aperture in the Fourier plane of each objective lens. Lateral displacement of these limiting apertures away from the optical axis of an objective lens sets the projection angle, for which the angular deviation from the central optical axis increases with increasing displacement, as illustrated in figure 1(a). Thus, the use of two apertures either side of the optical axis of the objective lens, defines respective principal rays on which two separate tube lenses can be centred to enable two projection images to be recorded on adjacent regions of a single camera sensor.

**Figure 1:**
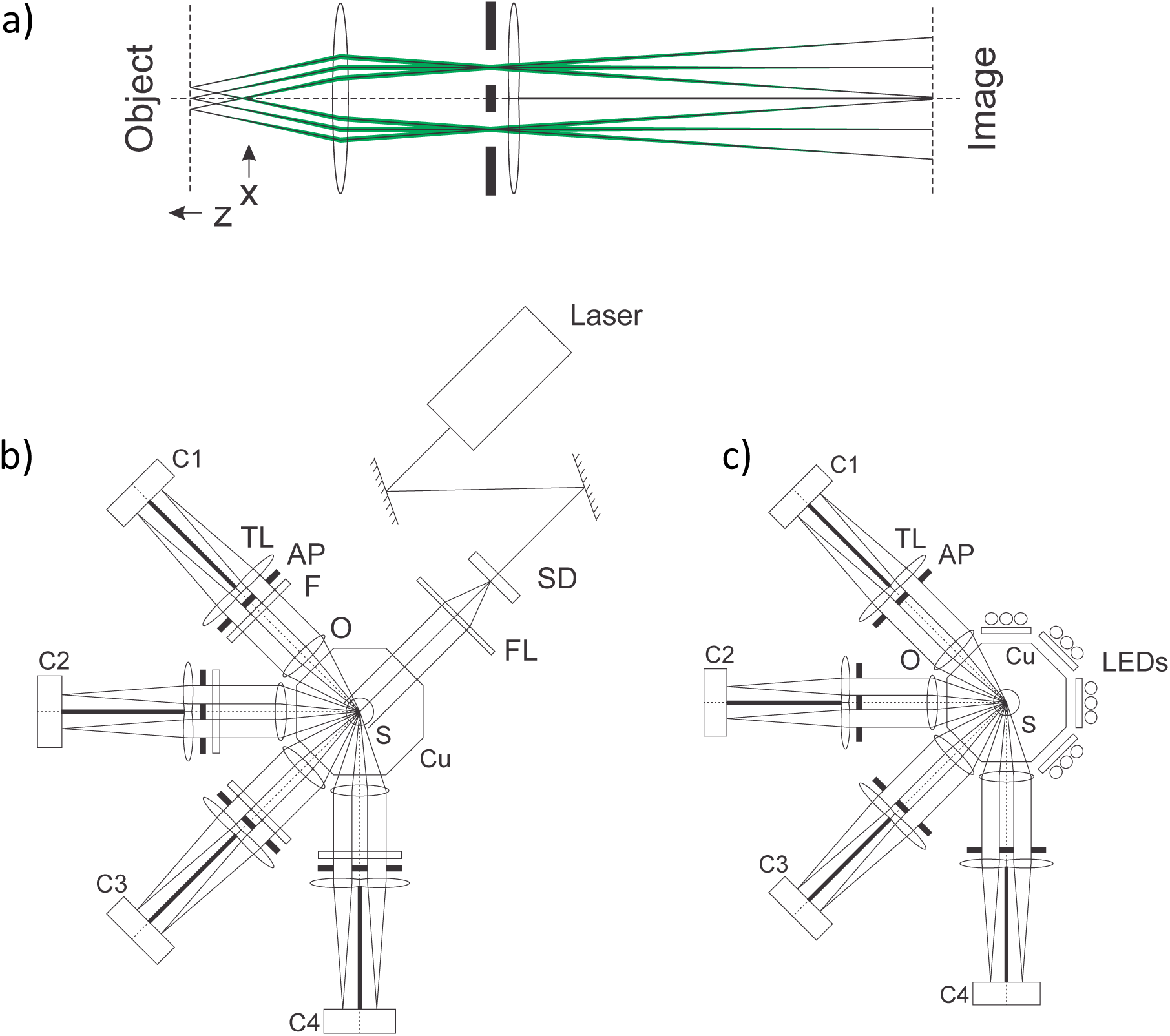
(a) A multiplexed imaging channel producing two images on a single camera. The two images of the object are at a relative angle of 3.8°, defined by aperture pinholes (AP) and composite tube lenses (TL) centred behind them. (b) Fluorescence and (c) transmitted light modalities of the multiplex multichannel single-shot OPT system. Time-averaged laser excitation is provided by a spinning diffuser (SD) and Fresnel lens (FL) combination and imaged through emission filters (F), while transmitted light is provided by white light LEDs with attached diffusers. The 8 images are recorded simultaneously on 4 synchronously triggered CMOS cameras (C1-4). Baffles (indicated as solid black line) are mounted between each camera and tube lens pair in order to minimise crosstalk.

For the results presented here, the apertures were set to fix the imaging NA to 0.028, corresponding to a depth of field of 1 mm, which was matched to the horizontal field of view of the system (limited by the size of the camera chip and the magnification). This ensured that a sample smaller than 1 mm was entirely within the depth of field. The corresponding aperture centre separation was 3.6 mm, yielding an angular separation of 3.8° between these projection images. The centres of the 125 mm focal length tube lenses were aligned to the centres of their respective apertures, which required truncating two doublet lenses [32-492 Edmund Optics] by cutting away their edges so only the centre regions of the lenses were used. This approach has previously been used for differential phase contrast microscopy^27^. To maximise the horizontal field of view (FOV), the tube lens lateral positions were set to locate the centre of images a quarter width from the camera sensor edges. To minimise crosstalk between multiplexed projection images, the tube lenses were placed directly behind the apertures, as depicted in Figure 1(a,b,c). As a result, each imaging arm was not image-space telecentric, although object-space telecentricity was preserved. Baffles were placed between the tube lenses in each imaging arm to prevent crosstalk from stray light.

The angular separation of each pair of multiplexed projection image arms was set to 45°, covering 135° on one side of the sample. This configuration was found to be optimal by acquiring a fully sampled 400-projections OPT dataset of a 3D bead sample (which was rotated using a motorised stage) and subsampling this data set to explore the performance of different undersampling strategies. This entailed qualitatively inspecting the resulting images (reconstructed using the iterative TwIST reconstruction method^5^) for visual quality and quantitatively assessing them by calculating the self-similarity image metric (SSIM) with the fully sampled OPT image reconstruction.

Figure 1(b,c) shows a schematic of the optical set-up of this single-shot OPT system. The sample was immersed in an index matching fluid contained in a custom-made octagonal cuvette, with the optical axes of the pairs of projection images at an angle of 1.9° either side of the normal to the cuvette surface. This small deviation from normal incidence had no measurable detrimental effect on the image quality due to the low NA of the imaging channels. For fluorescence OPT, 473nm excitation was provided by a diode laser (Cobolt Blues, Hubner Gmbh) that was directed through a spinning diffuser and collimated by a Fresnel lens to provide uniform wide-field illumination through a separate face of the cuvette. For transmitted light OPT, white LEDs were positioned on the opposite cuvette face of each pair of imaging channels. (fig.1b and c respectively). For the set-up and optimisation of the instrument, a stepper motor rotation stage (NM11AS-T4, Laser 2000 (UK) Ltd) was employed, but this was not used for the single-shot OPT acquisitions.

### Alignment & Registration

The correct alignment of a standard OPT set-up (where the object is rotated) requires that the rotation axis is centred in the field of view without tilt. This ensures that projection views are back-projected across reconstruction space in the correct position, and the object is reconstructed without misalignment artefact^28^. This alignment can be performed by comparing views of a test object before and after a rotation of 180°.

For the single-shot OPT system, we require the y-axes of all four pairs of imaging arms to be parallel to the y-axis in the object space, y_0_, as depicted in figure 2. This is equivalent to requiring that the x-z planes 1-4 are coplanar. If this condition is met, then an object can be correctly reconstructed from projection images recorded by each independent imaging arm, and is equivalent to aligning the rotation axis in standard OPT. This condition can be achieved by separately aligning each imaging arm using the standard OPT method (if a sample rotation stage is available). As each arm is aligned to the common vertical axis, it ensures co-registration between all the imaging channels. It is possible to align the system without using a rotation stage however, since a test object with a known 3D structure can be used to calculate the position and viewing angle of an imaging system^29^.

**Figure 2:**
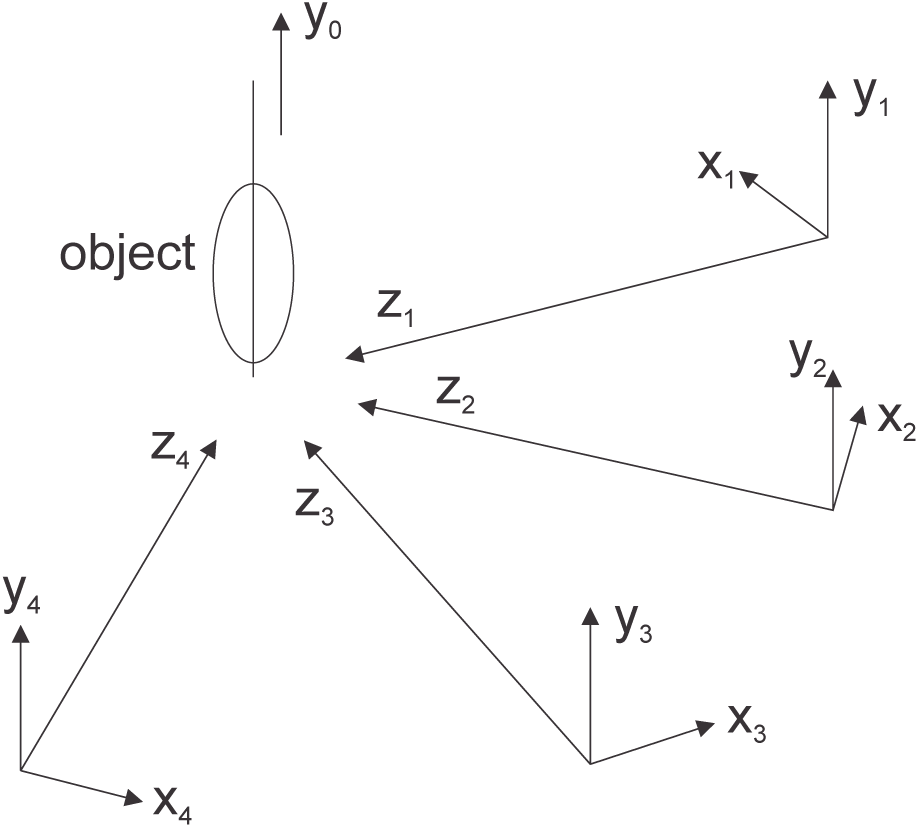
Demonstration of coordinate axis in ss-OPT. Global y-axis y0 indicates the vertical axis in object space. z1-4 are optical axes for each optical arm respectively. x1-4 and y1-4 are the horizontal and vertical axes in each camera plane. y0-4 should be parallel for proper system alignment.

**Figure 3:**
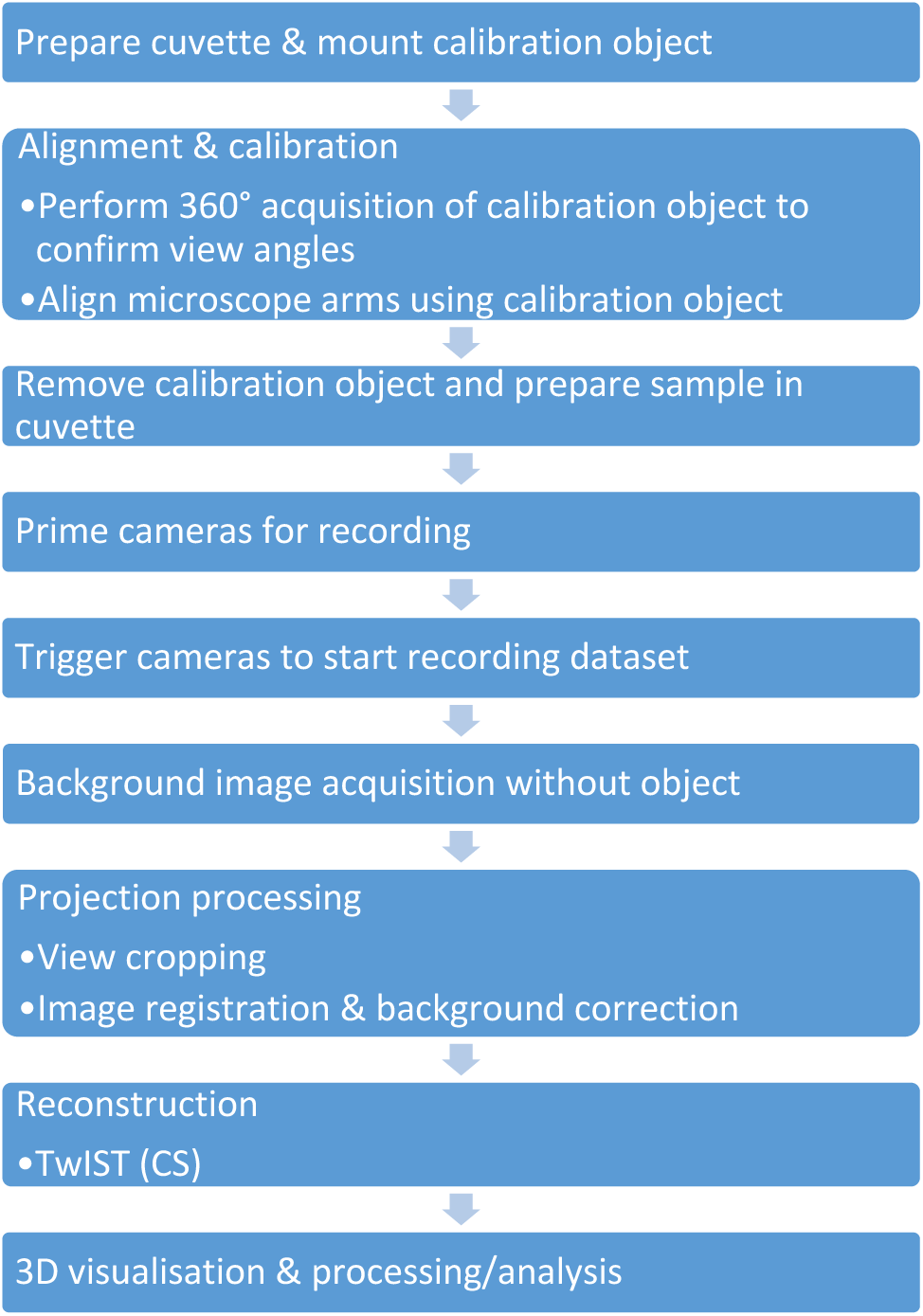
Data processing flowchart for the system

Co-registration of the four imaging arms was performed as part of the initial set-up of the single shot OPT image system. In the work reported here, each imaging arm was aligned in turn as outlined above. An OPT acquisition of a test object was then taken, recording 1280 projections over 360°. This dataset was used to measure the angular separation between projection views and cameras and was also used to confirm the registration of each imaging arm, and co-registration between all 4 arms. Based on this test, cropping of each projection view was applied to separate the two views on each camera and thus all 8 views were co-registered.

However, if such a fully sampled OPT dataset is not available, knowledge of the angular separation between cameras is sufficient (providing that all optical axes z^1-4^ are coplanar) for registration of ss-OPT image data utilising a single reference point in all projection views. As projections are parallel (i.e., the object lies within the telecentric depth of field), any common point at the intersection of the imaging arms can form the centre of the reconstruction space. By applying appropriate shifts to each camera image based on geometrical consideration of the angular separation between cameras, the imaging arms can be co-registered to enable the ss-OPT image data to be reconstructed.

### Volumetric reconstruction

Considering the imaging systems to be diffraction limited with a spatial resolution of 9.5 μm (for 520 nm light), the horizontal FOV and imaging NA resulted in ~ 84 resolution elements across a projection image (0.8 mm across). Therefore, ~ 262 projection views would be required for Nyquist sampling of an object filling the field of view (to enable artefact-free Fourier back-projection). Since the ss-OPT system acquired only 8 projection images, it was undersampled by a factor of 33x. We have previously shown that the TwIST algorithm can reconstruct undersampled OPT data (with as few as 20 projection images^5^) acquired with conventional OPT instruments and here we extend this approach, applying TwIST to retrieve the volumetric information from a ss-OPT dataset containing only 8 projection views. We note that other reconstruction methods may be used, such as the training of a convolutional neural network to remove the streak artefacts from the reconstructed volumetric data^25^.

### ss-OPT data acquisition

Figure 2 shows the full acquisition workflow for a single-shot OPT acquisition with the system. Four cost-effective CMOS cameras (Chameleon 3 CM3-U3-31S4M, Teledyne FLIR) with hardware triggering were used to capture images simultaneously. The hardware trigger signal was provided by an Arduino, which controlled the trigger pulse frequency, and a hardware switch was wired to start and stop the trigger pulse train.

A MATLAB programme was used for camera control during the OPT system registration and calibration, but ss-OPT acquisitions were recorded using SpinView, the software provided with the CMOS cameras. Single shot projection images were streamed onto a desktop computer and stored in a RAM buffer, which was continuously written to disk. An NVMe SSD drive was installed in the PC to maximise data write speed. If the disk write speed is too low, the maximum acquisition length will be limited by the RAM buffer capacity.

For transmission acquisitions, background images were recorded without the object present for each camera view, both with and without illumination. These were then used to correct each projection view and account for any intensity variations.

Projection image registration, TwIST reconstruction and binary masking were performed in MATLAB using custom scripts. Volume visualisation and rendering was performed using 3DScript^30^, an open source plugin for ImageJ, with macros written to render multiple timepoints from a ss-OPT dataset. Rendering settings may be fine-tuned to suppress any remaining streak artefacts and background signal in the final volumetric renders. Adobe Photoshop was used to colour code and label some renders.

## Results

Figure 4 addresses the reconstruction of undersampled OPT data and compares the reconstruction performance of the TwIST algorithm and filtered back projection, for both 200 projections (approximately fully sampled) and 8 projections. The test dataset is the fluorescence labelled vasculature of a 3 d.p.f. (days post fertilisation) fli:GFP zebrafish embryo. Panels 4(a-c) show single reconstructed slices from the dataset using these reconstruction methods, and 4(d-f) present a maximum intensity projection images through a section of the volumetric reconstruction. This OPT data was generated by recording a fully sampled (400 projection) dataset and subsampling the relevant projections. When using FBP with only 8 projections 4(c,f), the resulting reconstructed image is corrupted by streak artefacts, reducing contrast and making it impossible to resolve features. However, the image reconstructed from only 8 projection images using TwIST is clearly able to resolve the vascular structure 4(b,e). This supports our undersampled single-shot approach to OPT.

**Figure 5:**
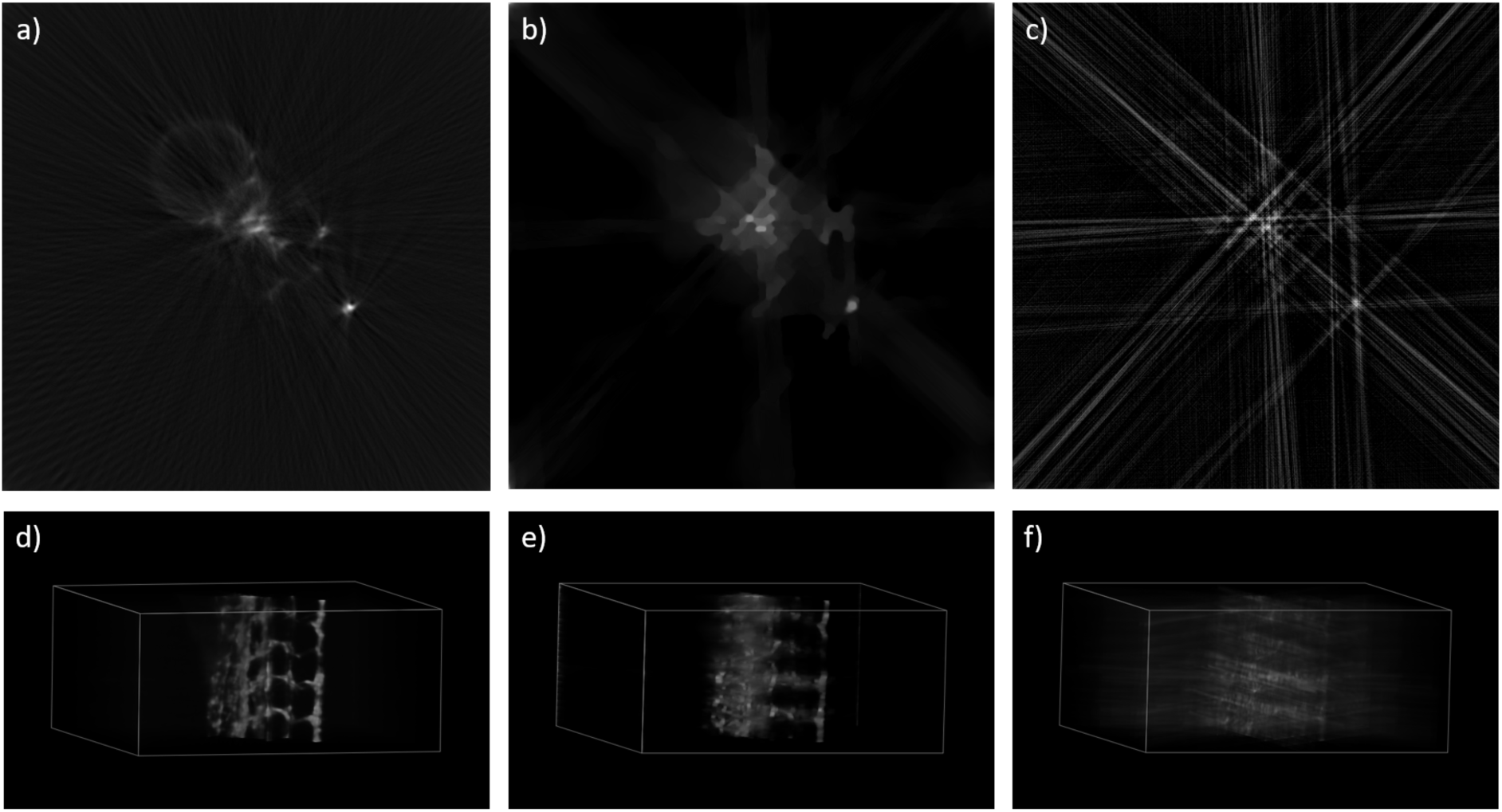
Comparison of FBP reconstruction with 200 projections (a,d), TwIST with 8 projections (b,e) and FBP with 8 projections (c,f). Dataset is fluorescence labelled vasculature in a 3d.p.f. fli:GFP zebrafish embryo. 8 projection dataset was generated from undersampling the full set of projection images. (a-c) Single slices through the reconstructed dataset. (d-e) renders of the reconstructed volumetric dataset, scale bars = 100µm. Intensity has been visually matched across the reconstruction methods to indicate features and detail present in the reconstructed volumes.

The peak-signal-to-noise (PSNR) values for the single reconstructed slices presented in fig. 4(b,c) when compared to the reference slice in fig. 4a are 18.39 dB for the TwIST reconstruction from 8 projections (fig 4b), and 13.18 dB for the FBP reconstruction from 8 projections (fig. 4c). As expected, the TwIST reconstruction results in a higher PSNR value, suggesting a closer representation of the ground truth is obtained when the TwIST algorithm is used from only 8 projections.

The first application of the single-shot OPT technique presented here is the (absorption) imaging of black ink flowing in water. The system was set up with transmission illumination, as shown in Fig. 1(c). The black ink was injected into the water-filled cuvette using a needle (also visible in the reconstructed images). The projection views were recorded every 62 ms, resulting in a ~ 16 volumes/second 3D time-lapse image dataset after reconstruction with the TwIST algorithm. Figure 5 presents four timepoints from this volumetric reconstruction. The intensity has been inverted to aid visualisation (i.e. brighter parts of the reconstruction correspond to greater absorption in the sample). The reconstruction has been colour-coded to encode the depth within the reconstructed volume for representation in a 2D figure. Supplementary videos 1 & 2 demonstrate the volumetric reconstruction over time, showing the shape and flow of the ink. This dataset demonstrates the ss-OPT technique’s potential for studies of fluid dynamics, including the capture of transient events, where contrast may be provided by either absorption or fluorescence.

**Figure 5:**
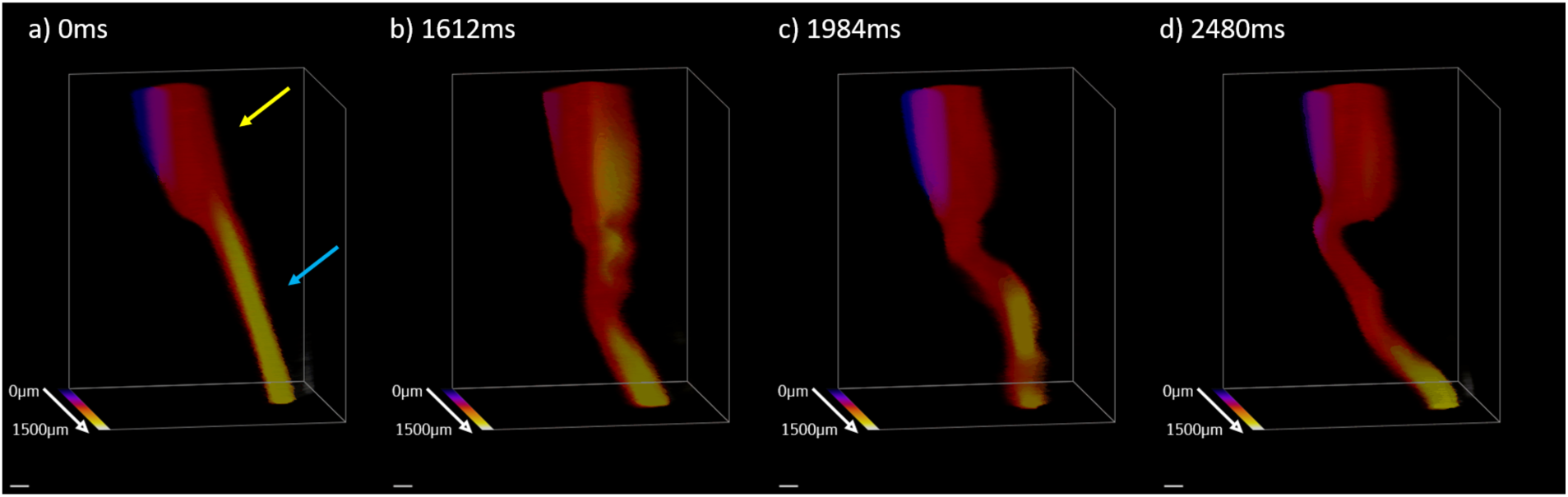
Reconstructions of a single-shot OPT acquisition of absorption due to black ink flowing in water, at 4 different timepoints from a continuous dataset: 0ms, 1612ms, 1984ms & 2480ms. The image intensity has been inverted from the original projection images for clarity. Dataset was recorded at ~ 16vol/sec, with a volume recorded every 62ms. Yellow arrow indicates the needle through which the ink (indicated by the blue arrow) was injected into the water. Colour represents depth within the reconstructed volume, as indicated by the colour bar. Scale bar = 100µm.

We now present the imaging of absorption (or attenuation coefficient) in a live biological sample using the ss-OPT method. The sample used was a 4 d.p.f. wild-type zebrafish embryo. Wild-type embryos were selected as their pigmentation provides contrast throughout their body, but scattering was still acceptably low due to the young age of the embryos. As the ss-OPT technique can image so rapidly, there was no need to anaesthetise the embryos before imaging. Thus, the zebrafish embryos were volumetrically imaged while awake and moving.

Figure 6 shows a single timepoint from a single-shot acquisition of a 4 d.p.f. zebrafish embryo as described above, with the volumetric reconstruction viewed from 4 angles. Projection views were recorded every 16 ms, with a camera integration time of 8 ms, resulting in a 62.5 volumes/second dataset, the limit of the cameras used. The embryo was mounted in a tube filled only with water, and the embryo was able to swim within the tube. The inner diameter of the mounting tube was 0.8 mm, chosen such that the sample would stay within the FOV of the system. The intensity values of the reconstructed have again been inverted to aid visualisation. Supplementary video 3 shows 1280 ms of the volumetric time lapse dataset, displayed from the four view angles seen in figure 6. It is possible to observe the beating of the heart in the reconstruction, and this is highlighted in supplementary video 4.

**Figure 6:**
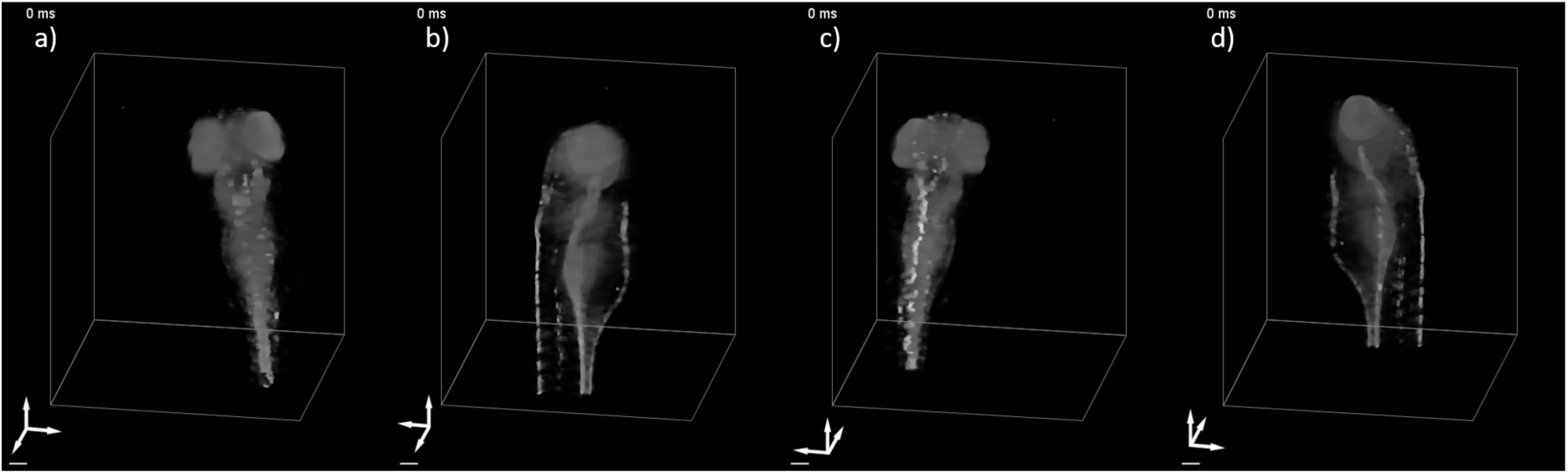
Four views of a single reconstructed volume from a single-shot OPT acquisition of the absorption in a 4d.p.f. wild-type zebrafish embryo. The image intensity has been inverted from the original projection images for clarity. The reconstructed volume has been rotated 90° between each view (a-d), as indicated by the axis labels. The zebrafish embryo was allowed to freely swim within the mounting tube, which was in turn immersed in the cuvette. The embryo was not placed under anaesthetic. The dataset was recorded at 62.5vol/sec, with projection views recorded every 16ms. Camera integration time = 8ms. Scale bar = 100µm.

Figure 7 shows 6 timepoints from later in the same ss-OPT time-lapse acquisition featured in figure 6: here the tail of the embryo was in the imaging FOV and the mounting tube was set such that the tail moved through the reconstructed volume while the embryo was swimming. The depth within the reconstructed volume is colour coded. This ss-OPT time-lapse data is also presented in video 5.

**Figure 7:**
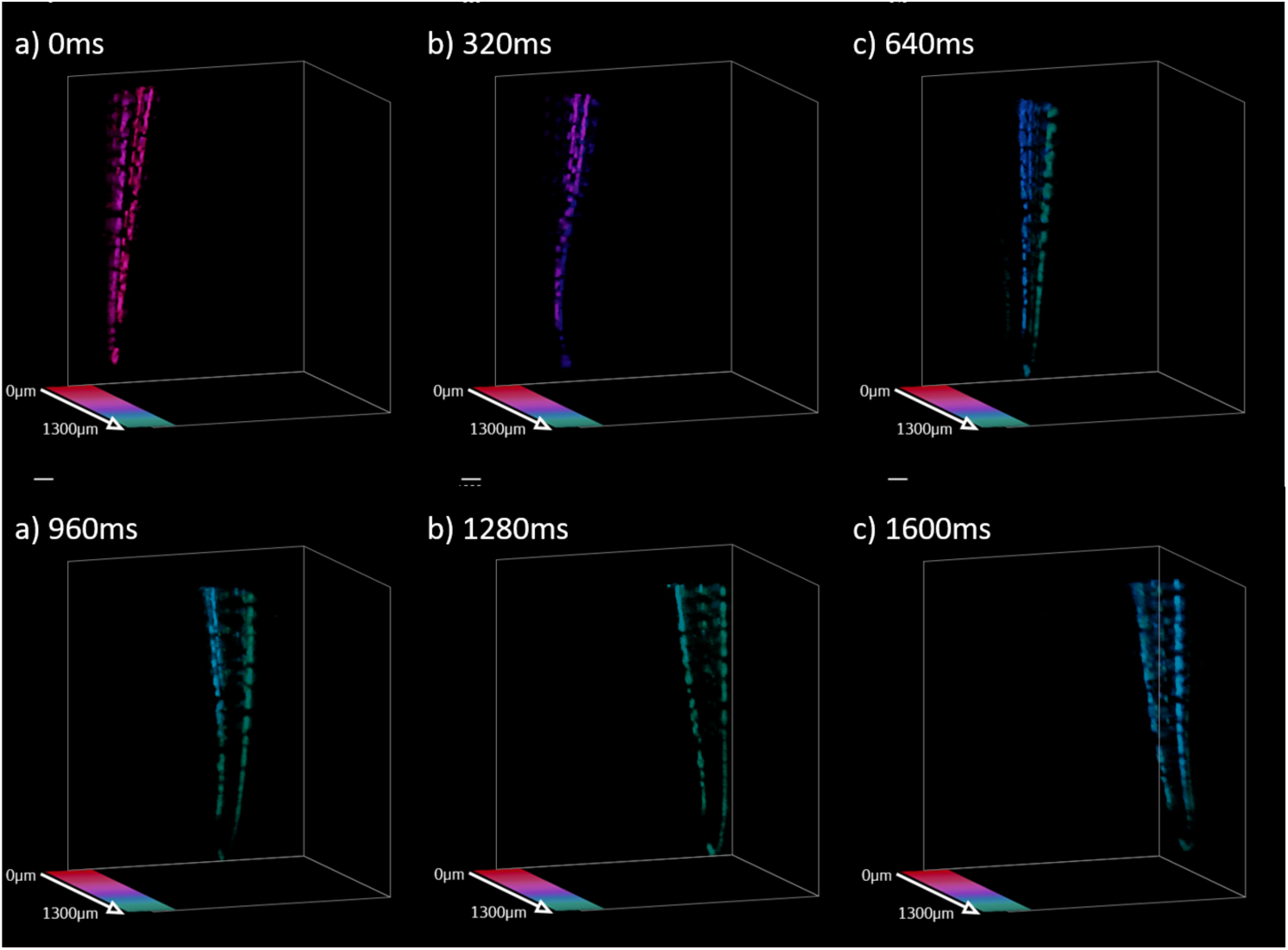
Six timepoints (0ms, 320ms, 640ms, 960ms, 1280ms, 1600ms) from the reconstruction of a single-shot OPT acquisition of the absorption in the tail of a 4d.p.f. wild-type zebrafish embryo, recorded at 62.5vol/sec. The image intensity has been inverted from the original projection images for clarity. The embryo was not placed under anaesthetic, and was allowed to swim freely within the 1mm inner diameter mounting tube. The mounting tube was rotated over the acquisition. Colour represents depth within the reconstructed volume, as indicated by the colour bar. Scale bar = 100µm.

A 4 d.p.f. zebrafish embryo was then mounted in a tube with a 5 mm inner diameter, allowing it to swim freely in any direction, into and out of the field of view of the system. This led to truncation of the object during the time-lapse acquisition and out of focus light was detected from parts of the sample along certain projection views. The camera integration time was reduced to 5 ms because the embryo was able to travel further in a fixed time window due to the increased space and this rapid movement of the embryo resulted in motion blur appearing in some projection images. The projections were again recorded at the maximum camera frame rate of 62.5 frames/second. Volumetric image data at three time points from the dataset are presented in Figure 8, with colour encoding the depth in the reconstructed volume. It can be seen that the embryo is able to freely orient itself within the mounting tube and leave the field of view in any direction. Videos 6 & 7 present 80 timepoints from this single-shot OPT time-lapse dataset.

**Figure 8:**
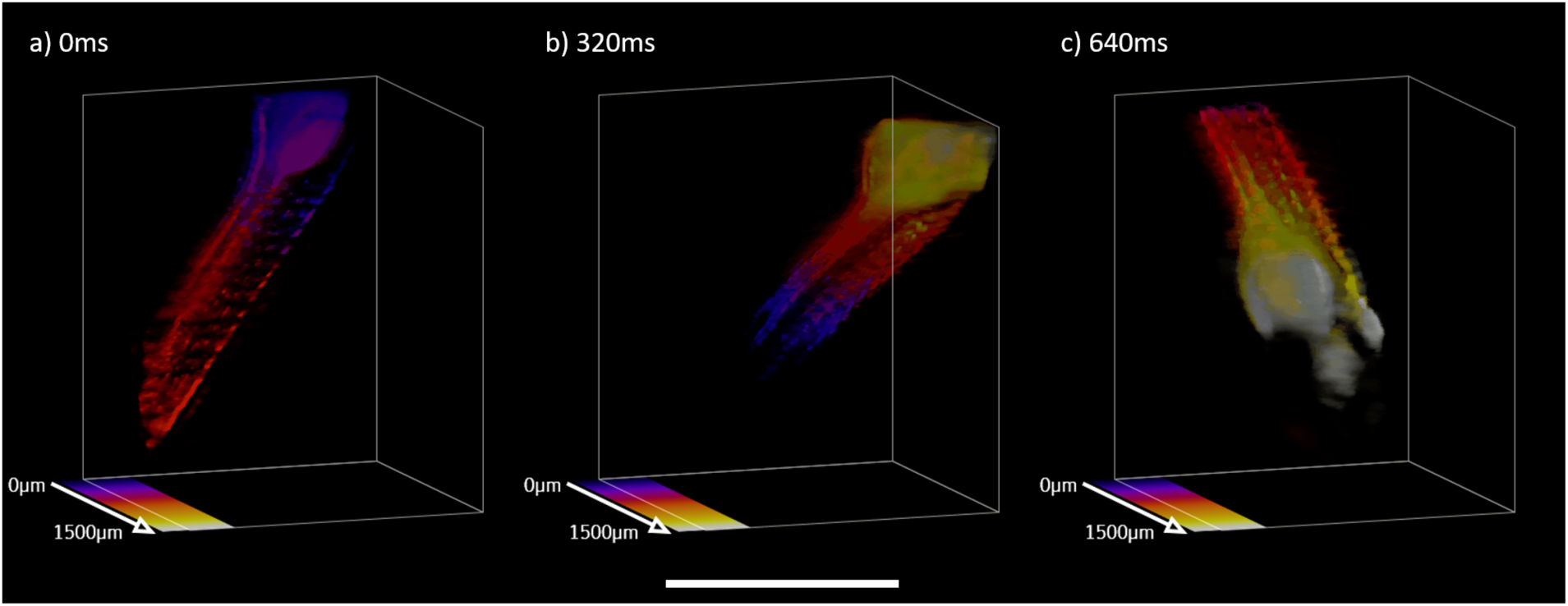
Reconstruction of a single-shot OPT absorption acquisition of a freely swimming 4d.p.f. wild-type zebrafish embryo, recorded at 62.5vol/sec. Intensity has been inverted from the original projection images for clarity. The embryo was not placed under anaesthetic and was allowed to freely swim within the 1cm inner diameter mounting tube, and so was free to leave the field of view. Projection views were recorded every 16ms, and volumetric reconstructions for views recorded at 64ms (a), 320ms (b) and 544ms (c) are presented here. Colour indicates depth within the reconstructed volume, as indicated by the colour bars. Scale bar = 1000µm.

Figure 9 presents 12 timepoints from the same volumetric dataset. Each frame is labelled in the render with a unique colour, as indicated in the colour-scale bar. Multiple volumes are presented simultaneously in each panel (as indicated by the number next to each render) in order to illustrate the motion of the embryo over time. Some volumes (e.g., at timepoints 3 and 7) show visible burring in the reconstruction due to the rapid motion of the fish during the 5 ms integration of the cameras. Video 8 presents the data depicted in figure 9 in motion. Video 9 presents a longer excerpt of the ss-OPT dataset, showing 5 reconstructed volumetric images from different timepoints in parallel. Previous and subsequent frames are colour coded in order to demonstrate the movement of the embryo over time.

**Figure 9:**
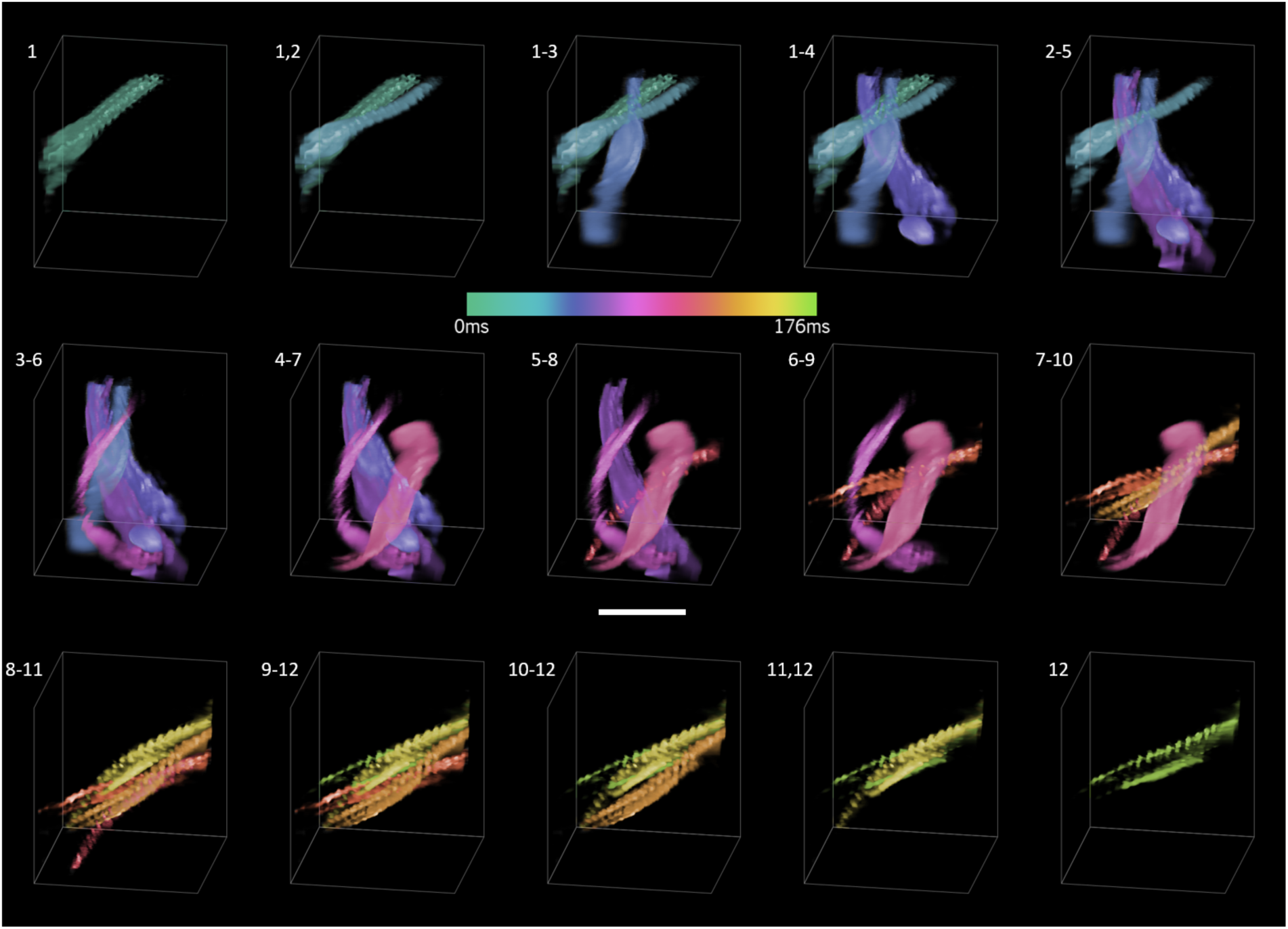
Reconstruction of a single-shot OPT absorption acquisition of a freely swimming 4d.p.f. wild-type zebrafish embryo, recorded at 62.5vol/sec. Intensity has been inverted from the original projection images for clarity. The embryo was not placed under anaesthetic and was allowed to freely swim within the 1cm inner diameter mounting tube, and so was free to leave the field of view. Projection views were recorded every 16ms. Up to 4 volumes are shown simultaneously, as indicated by the frame number next to each panel, in order to demonstrate the motion of the embryo. Each rendered timepoint has been assigned a unique colour, as shown in the colour bar. As the fish was freely moving during the acquisition, some projection views were blurred due to fast movement of the embryo while the camera sensors were integrating. This was minimised by reducing the integration time, however due to the distance moved by the sample over this 5ms timescale, some blurring is still apparent. Scale bar = 1000µm.

## Conclusions

We have here presented a proof-of-principle ss-OPT system that simultaneous acquires 8 projection images using four low-cost CMOS cameras and 4x microscope objective lenses. Please see Supplementary Information for a complete list of components for this system. The optical components of this ss-OPT system may be adapted to provide different performance in terms of magnification, field of view, imaging rate and image quality, e.g., by changing the imaging objective lens, tube lens and/or camera (sensor size, quantum efficiency, noise, frame rate) and by changing the number of imaging arms. It would also be possible to combine fluorescence and transmitted light imaging. The capability for rapid volumetric imaging – primarily limited only by the frame rate of the camera or the signal level – may make this a useful approach for 3D flow cytometry and/or sorting, e.g. of small organisms or 3D organoids, and may also be applied to study fluid dynamics. The ability to implement an OPT system without the need for sample rotation may also have applications in situations here sample rotation is not practical, such as developmental biology or plant imaging.

Although the single-shot datasets presented here measure absorption, we have demonstrated that our compressed sensing approach is valid for fluorescence acquisitions (fig. 4), and measurement of fluorescence signals in dynamic samples will be the subject of future work.

As well as improving the single-shot image quality by employing cameras with superior signal-to-noise properties or by increasing the number of imaging arms, it may also be possible to improve the image quality through advances in compressed sensing algorithms or in deep learning reconstruction methods. We note that CNN reconstruction provides faster data analysis once the training model is established while the iterative compressed sensing approach (e.g., TwIST) can be used for any sample and may be more robust with respect to changes in sample illumination or sample motion during the single-shot image acquisition time. This will be the subject of future work.

## Acknowledgements

We are grateful to Anna Rydlova and Margaret Dallman, Department of Life Sciences at Imperial College London, for providing the zebrafish embryos imaged during this work. The custom components in the instrument reported here were co-designed and fabricated by Simon Johnson, Martin Kehoe and John Murphy, Optics instrumentation facility in the Department of Physics at Imperial College London. We also gratefully acknowledge the donation of a Tesla K40 GPU by NVIDIA (https://registration.nvidia.com/ahr.aspx).

This work was supported in part by the Engineering and Physical Sciences Research Council (EPSRC, EP/V048996/1). Sunil Kumar is supported by funding from the Francis Crick Institute, which receives its core funding from Cancer Research UK (FC001999), the UK Medical Research Council (FC001999) and the Wellcome Trust (FC001999). Connor Darling acknowledges a PhD studentship from the EPSRC National Productivity Investment Fund with Cairn Research Ltd and Samuel Davis acknowledges a PhD studentship from EPSRC.

For the purpose of Open Access, the author has applied a CC BY public copyright licence to this manuscript.

