## Supplementary Material for "Single-Shot Optical Projection tomography for high-speed volumetric imaging"

**List of Supplemental Videos**

*Video 1: Reconstruction of a single-shot OPT acquisition of absorption due to black ink flowing in water. Dataset was recorded at ~16vol/sec, with a volume recorded every 62ms. Colour represents depth within the reconstructed volume, as indicated by the colour bar. Scale bar = 100µm.*

*Video 2: Reconstruction of a single-shot OPT acquisition of absorption due to black ink flowing in water. Dataset was recorded at ~16vol/sec, with a volume recorded every 62ms. Rendering viewpoint is rotated over the video in software in order to aid visualisation of the 3D dataset on a 2D monitor. Scale bar = 100µm.*

*Video 3: 80 volumes from a single-shot OPT acquisition of a 4d.p.f. zebrafish recorded at 62.5vol/sec. Projection views were recorded every 16ms, as indicated by the time counter in the top right. The embryo was not placed under anaesthetic and allowed to freely swim within the mounting tube. The 80 volumes from the dataset are replayed from 4 reconstruction viewpoints, controlled in the rendering software (3DScript). Video playback is at 20fps, resulting in slowing of the video to 313% original duration. Scale bar = 100µm*

*Video 4: Left: Single render view of reconstruction from video 7, in which the pumping of the embryo heart can be observed. Right: Inverted intensity single camera projection view of same time points.*

*Video 5: Reconstruction of a single-shot OPT acquisition of the absorption in the tail of a 4d.p.f. wild-type zebrafish embryo, recorded at 62.5vol/sec. Intensity has been inverted from the original projection images for clarity. The embryo was not placed under anaesthetic, and was allowed to swim freely within the 1mm inner diameter mounting tube. The mounting tube was rotated over the acquisition. Colour represents depth within the reconstructed volume, as indicated by the colour bar. Scale bar = 100µm.*

*Video 6: Reconstruction of a single-shot OPT absorption acquisition of a freely swimming 4d.p.f. wild-type zebrafish embryo, recorded at 62.5vol/sec. Intensity has been inverted from the original projection images for clarity. The embryo was not placed under anaesthetic and was allowed to freely swim within the 1cm inner diameter mounting tube, and so was free to leave the field of view. Projection views were recorded every 16ms, as indicated by the volume time count in the upper left. Video playback is at 30fps, resulting in slowing of the video to 208% original duration. Scale bar = 100µm.*

*Video 7: Reconstruction of a single-shot OPT absorption acquisition of a freely swimming 4d.p.f. wild-type zebrafish embryo, recorded at 62.5vol/sec. Intensity has been inverted from the original projection images for clarity. The embryo was not placed under anaesthetic and was allowed to freely swim within the 1cm inner diameter mounting tube, and so was free to leave the field of view. Projection views were recorded every 16ms, as indicated by the volume time count in the upper left. Video playback is at 20fps, resulting in slowing of the video to 313% original duration. Colour indicates depth within the reconstructed volume, as indicated by the colour bars. Scale bar = 100µm.*

*Video 8: 12 volume excerpt of single-shot acquisition of a freely swimming 4d.p.f. wild type zebrafish embryo, recorded at 62.5vol/sec. The volumetric reconstruction of each time point has been assigned a unique colour, as indicated by the colour bar (top). Up to 4 volumes are displayed simultaneously in order to demonstrate motion of the embryo over time. Scale bar = 100µm.*

*Video 9: Single-shot OPT acquisition of a freely swimming 4d.p.f. wild-type zebrafish embryo, recorded at 62.5vol/sec. Projection views were recorded every 16ms. Volumetric reconstructions from 5 timepoints are presented simultaneously in each video frame here, with the current frame in pink, and two previous and subsequent frames colour coded as shown in the colour bar (top). Scale bar = 100µm.*

**List of components**

- Custom 8-walled glass cuvette (octagonal cross-section).
- Zaber NM11AS-T4 Stepper Motor[1], with A-MCA controller[2]
- Cobolt Blues 473nm 50mW diode laser[3]
- Workstation with Nvidia K40c GPU, with 12GB of GPU memory, for reconstruction.
- Arduino Uno[4], for hardware triggering cameras
- Per Imaging arm:
    - AmScope 4x Plan Fluor Objective Lens[5]
    - Edmund Optics 25mm diameter 125mm focal length achromatic doublet[6] x2, truncated
    - FLIR Chameleon3 CM3-U3-31S4M-CS CMOS Camera[7]
    - Thorlabs 30mm cage components[8]
    - Custom aperture plate, with 2 aperture pinholes drilled, centres 3.6mm apart
    - LED array with built in diffuser

### References


1. NM11AS-T4 Specifications - Zaber. https://www.zaber.com/products/stepper-motors/NM/details/NM11AS-T4?sr=.

2. A-MCA Specifications - Zaber. https://www.zaber.com/products/controllers-joysticks/A-MCA/details/A-MCA?sr=.

3. Cobolt 04-01 Series - HÜBNER Photonics - Lasers & THz systems. https://hubner-photonics.com/products/lasers/single-frequency-lasers/04-01-series/.

4. Arduino. Arduino Uno Rev3: Documentation. https://store.arduino.cc/arduino-uno-rev3 (2018).

5. Amscope. 4X Infinity-corrected Plan Fluor Objective Lens. https://www.amscope.com/accessories/objective/4x-infinity-corrected-plan-fluor-objective-lens.html.

6. 25mm Dia. x 125mm FL, MgF2 Coated, Achromatic Doublet Lens. https://www.edmundoptics.co.uk/p/25mm-dia-x-125mm-fl-mgfsub2sub-coated-achromatic-doublet-lens/2376/.

7. FLIR Machine Vision. Chameleon3 3.2 MP Mono USB3 Vision (Sony IMX265). https://www.ptgrey.com/chameleon3-32-mp-mono-usb3-vision-sonyimx265.

8. 30 mm Cage Components - Thorlabs. https://www.thorlabs.com/navigation.cfm?guide_id=2004.
